# Differences in anti-viral immune responses in individuals of Indian and European origin: relevance for the COVID-19 pandemic

**DOI:** 10.1101/2022.08.30.505791

**Authors:** Büsranur Geckin, Martijn Zoodsma, Gizem Kilic, Priya A. Debisarun, Srabanti Rakshit, Vasista Adiga, Asma Ahmed, Chaitra Parthiban, Nirutha Chetan Kumar, George D’Souza, Marijke P Baltissen, Joost H A Martens, Jorge Dominguez Andres, Yang Li, Annapurna Vyakarnam, Mihai G Netea

## Abstract

During the COVID-19 pandemic, large differences in susceptibility and mortality due to SARS-CoV-2 infection have been reported between populations in Europe and South Asia. While both host and environmental factors (including BCG vaccination) have been proposed to explain this, the potential biological substrate of these differences is unknown. We purified peripheral blood mononuclear cells from individuals living in India and the Netherlands at baseline and 10-12 weeks after BCG vaccination. We compared chromatin accessibility between the two populations at baseline, as well as gene transcription profiles and cytokine production capacities upon viral stimulation with influenza and SARS-CoV-2. The chromatin accessibility of genes important for adaptive immunity was higher in Indians compared to Europeans, while the latter had more accessible chromatin regions in genes of the innate immune system. At the transcriptional level, we observed that Indian volunteers displayed a more tolerant immune response to viral stimulation, in contrast to a more exaggerated response in Europeans. BCG vaccination strengthened the tolerance program in Indians, but not in Europeans. These differences may partly explain the different impact of COVID-19 on the two populations.

## Introduction

The immune response to pathogens is elicited and regulated through a series of pathways shaped by genetic and environmental factors. While the genetic component is constant, the environmental factors continuously change during an individual’s life course and their cumulative effect regulates the immune response (1). These environmental factors include lifestyle, diet, vaccinations, infections, and treatments. By extension, this leads to a great variation in response to pathogens at the individual level. Subsequently, such heterogeneity at the individual level translates into distinct responses in various populations, leading to differences in disease outcome (2,3).

During the COVID-19 pandemic, striking observations were made regarding differences in the impact and outcome of SARS-CoV-2 infection between populations. It has been reported that developing countries such as India and countries in sub-Saharan Africa had significantly lower mortality rates compared to populations in the developed nations of Western Europe and North America (4), despite possible biases due to under-reporting. Several mechanisms have been suggested to play a role, such as cross-reactive humoral and cellular lymphocyte responses to other coronaviruses or virus-virus interactions: a study examined serological cross-reactivity of COVID-19-naïve people from sub-Saharan Africa against SARS-CoV-2, which showed a high prevalence of pre-existing serological cross-reactivity to the novel virus (5). Another hypothesis proposed an increased level of natural resistance to explain, at least partly, the inter-populational variation in response to SARS-CoV-2, resistance that can be induced by local infectious pressure (6). Little is known, however, regarding the potential differential regulation of immune responses to SARS-CoV-2 or other viruses in various populations.

Considering these observations relevant for the COVID-19 pandemic, we aimed to understand the differences in anti-viral immune responses of individuals of Indian or European ancestry to the SARS-CoV-2 and influenza viruses. In addition, we studied whether these responses are affected by BCG vaccination, which has been recently suggested to be able to increase host defense against heterologous infections such as COVID-19.

## Results

### Chromatin accessibility profiles differ in Indian and European individuals

We obtained peripheral blood mononuclear cells (PBMCs) from ten Indian (6 men, 4 women, all between 20 and 30 years of age) and ten European (5 men, 5 women, also between 20 and 30 years of age) individuals before and 10-12 weeks after BCG vaccination (**Figure 1A**). First, we investigated the differences in chromatin accessibility in unstimulated PBMCs from Indian and European individuals before the BCG vaccination (during homeostasis). Differential peak analysis showed substantial differences between the two populations (**Figure 1B**). Subsequently, motif enrichment analysis on the differentially accessible peaks per population identified several over-represented transcription factor-binding motifs unique to each population (**Figure 1C**). In the European population, we found transcription factors CCAAT Enhancer Binding Protein Beta (CEBPB) and Hepatic Leukemia Factor (HLF) to be over-represented. CEBPB regulates the expression of genes that are involved in immune pathways and inflammatory responses (7). In the Indian population, multiple transcription factors of the RUNX family were significantly enriched, which are critical mediators of hematopoiesis [reviewed in (8)]. Furthermore, the transcription factor JunB is involved in a wide range of immunological processes, including macrophage activation (9) and regulatory T cell homeostasis (10).

**Figure 1.**
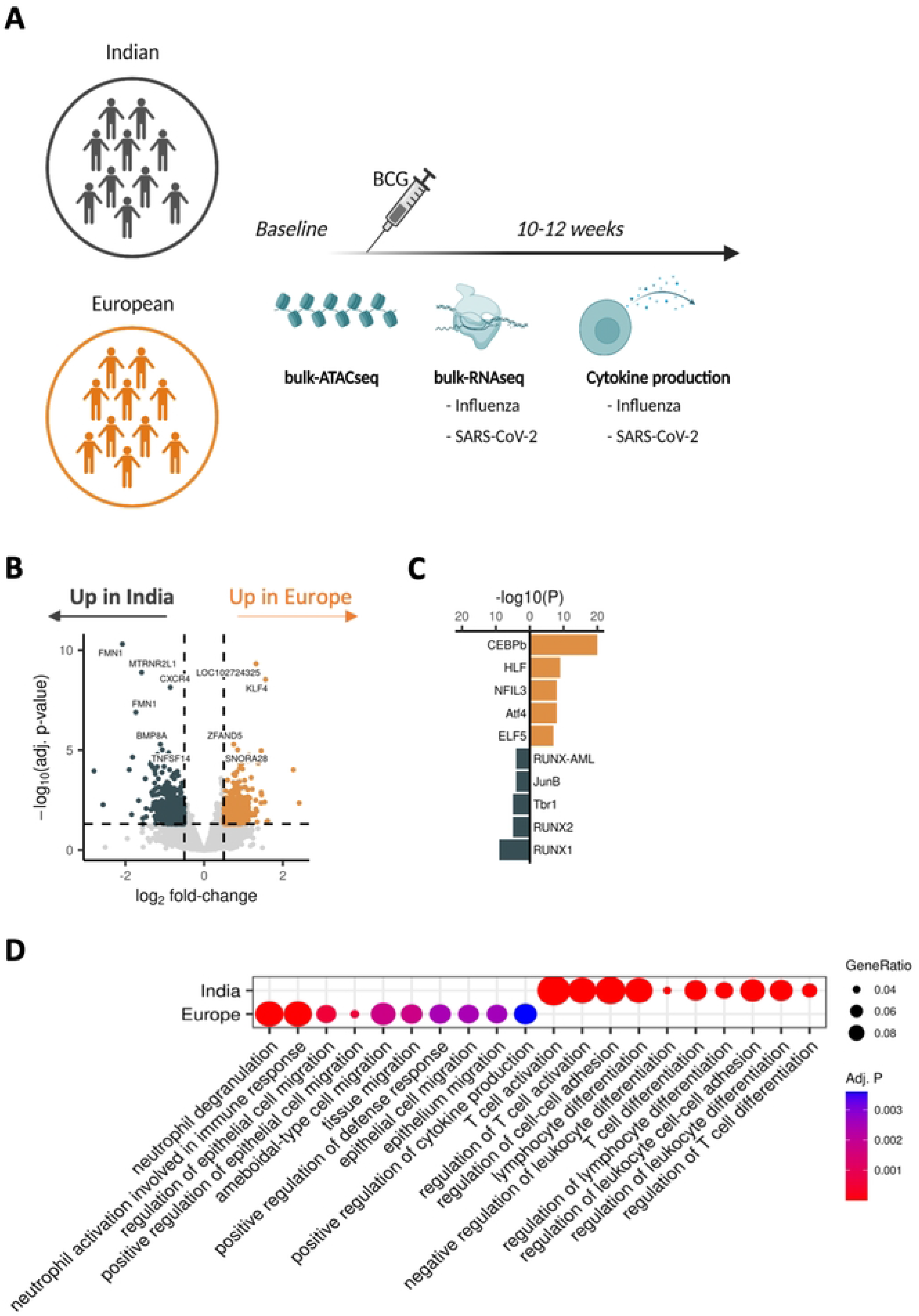
Chromatin accessibility profiles in Indian and European individuals demonstrates activity in different compartments of the immune system. **A)** Schematic overview of the study. **B)** Volcano plot showing the difference in chromatin accessibility per peak between European and Indian individuals at baseline (Absolute log2-transformed fold-change > 0.5, adj. P-value < 0.05). The most significant peaks are labeled based on their closest gene based on HOMER annotation. Golden peaks are upregulated in European individuals, while dark grey points are upregulated in Indian individuals. **C)** Barplot of the transcription factor motif enrichment among the differentially accessible peaks. **D)** Dot plot of the GO term enrichment on the genes linked to differentially accessible peaks.

Using gene set enrichment analysis of genes in close proximity to differentially accessible peaks (**Figure 1D**), we observed striking differences between the enriched pathways per population. Open peaks in the European population were more enriched in genes important for innate immune-related processes, whereas, open peaks in the Indian population were highly related to adaptive immunity and T cell immunity.

### Antiviral response is more regulated in Indian individuals compared to European individuals

We explored the transcriptional differences between Europeans and Indians in untreated conditions and upon stimulation with SARS-CoV-2 and Influenza, respectively. In untreated conditions, we found substantial differences between the populations at the transcriptional level (**Figure 2A**). Among the most significantly differentially expressed genes were FOSB and JUN, consistent with enrichment of the JunB transcription factor from our afore-mentioned analysis.

**Figure 2.**
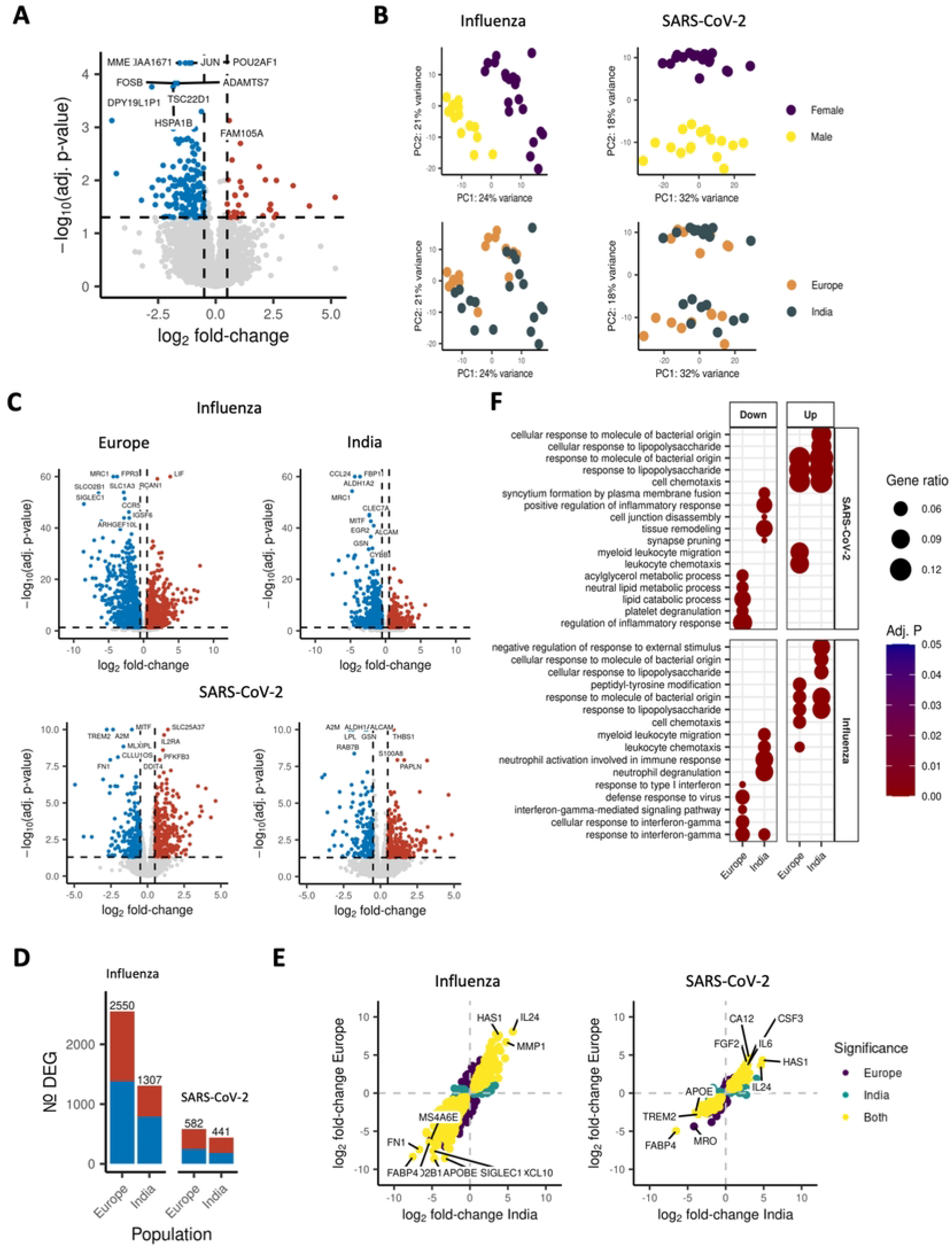
Transcriptional response to SARS-CoV-2 and Influenza viruses differs in magnitude between Europeans and Indians. **A)** Volcano plot of the baseline differences between European and Indian individuals (Absolute log2-transformed fold-change > 0.5, adj. P-value < 0.05). **B)** Principal component analysis (PCA) of the response to stimulation. Each point represents a sample. Samples included are those that were Influenza / SARS-CoV-2 stimulated. **C)** Volcano plots showing the response to Influenza and SARS-CoV-2 stimulation compared to RPMI-treated PBMCs. **D)** Barplot of the number of differentially expressed genes per stimulation and population. Blue indicates downregulation, red indicates upregulation. **E)** Consistency of the response to viral stimulation between the European and Indian populations. Each point represents a gene, plotted by the logFC per stimulation compared to RPMI in Indian and Europeans. **F)** Dotplot of GO term enrichment results of the differentially expressed genes per stimulation and population.

For PBMCs stimulated with either SARS-CoV-2 or Influenza, PCA-based dimensionality reduction revealed that the individual’s sex, and not their geographical origin, drove more strongly the variance between samples (**Figure 2B**). Overall, both viruses led to a considerable response at the transcriptional level **(Figure 2C**). Investigating the viral response in both populations revealed that Europeans were more responsive to viral stimuli compared to Indians, evidenced by a higher number of differentially expressed genes (**Figure 2D**) and a larger magnitude of up- and downregulation (**Figure 2E**). These effects were more visible for the Influenza virus compared to SARS-CoV-2. Finally, we asked which transcriptional modules were activated in response to viral stimulation, and how these differed between the populations. **Figure 2F** shows both shared and population-specific transcriptional modules significantly enriched upon stimulation. Europeans down-regulated more pathways related to type 1 interferons and interferon-gamma signalling in response to stimulation, while Indians more strongly down-regulated pathways related to myeloid cell migration and neutrophil activation, both important features of inflammation.

In agreement with the transcriptional data, cytokine production overall was higher in European individuals compared to the Indian group. Pro- and anti-inflammatory cytokines (IL-1Ra, IL-1b, IL6, TNFa) were all more strongly elevated in the European group after 24 hours of stimulation with the influenza virus. This difference reached significance (Two-sided Wilcoxon rank-sum test, p=0.001) in the IL-1Ra readout despite the small sample size. In contrast, response to SARS-CoV-2 showed large variation within the production of these cytokines. IL-1Ra production showed a clear difference as Indian individuals had lower production than European individuals. However, pro-inflammatory cytokines IL-1b, IL6, and TNFa production were higher in Indian individuals in response to SARS-CoV-2 **(Figure 3A**). Nevertheless, these differences were not statistically significant. Next, we checked T-cell derived cytokines to understand the antigen-specific responses better (**Figure 3B**). Production of the anti-viral cytokine IFNγ was higher in the European group, but statistical differences were reached only in response to SARS-CoV-2 stimulation. This pattern remained the same for IL-17: lower production in Indian individuals and higher in Europeans, but the difference was not significant. IL-10 showed no difference between the two populations. These data show that Indian individuals have an overall lower responsiveness to viral stimuli, i.e. influenza and SARS-CoV-2.

**Figure 3.**
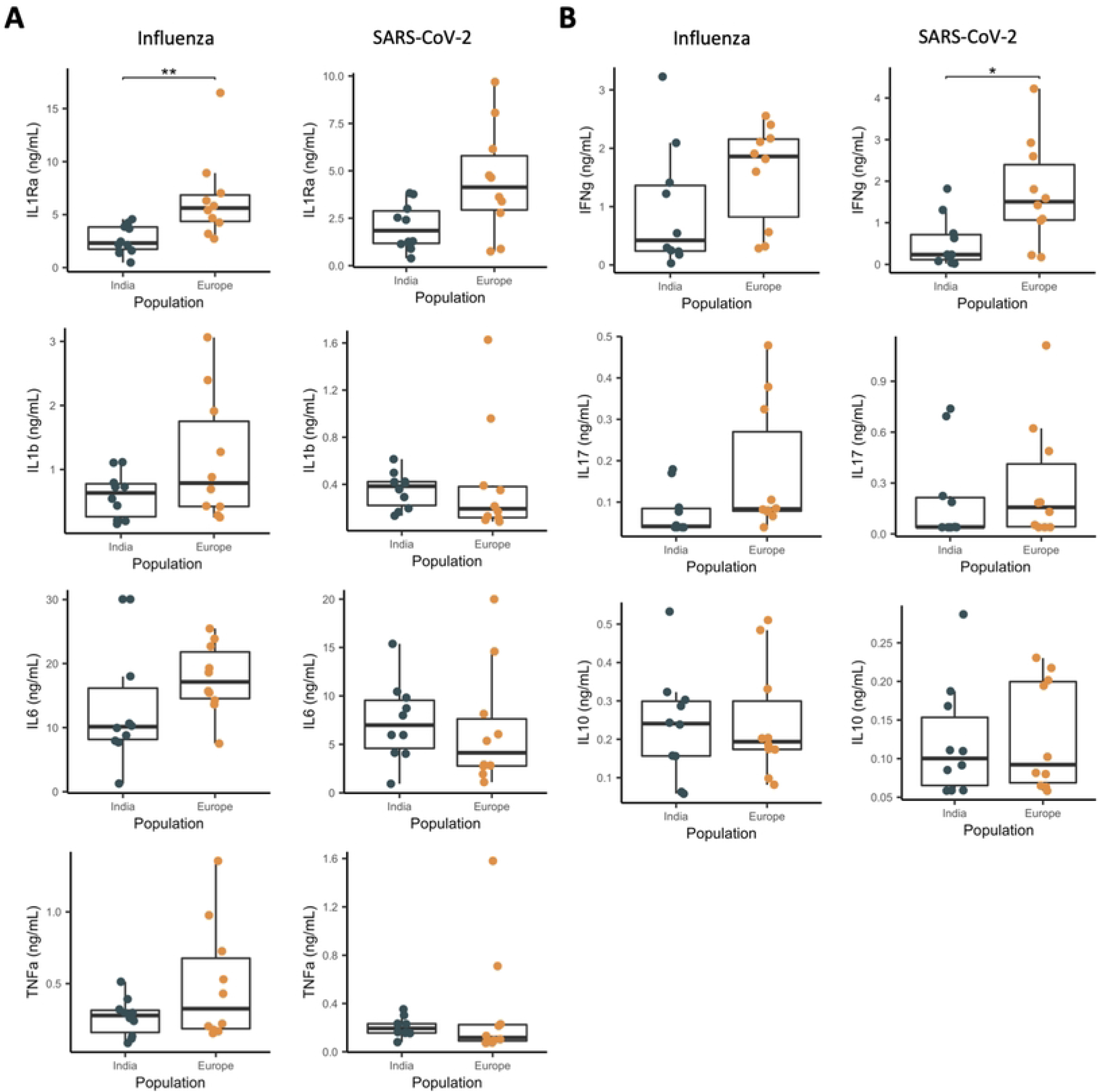
Cytokine production in Indian and European individuals. **A)** Pro-inflammatory cytokine production following 24 hours stimulation measured by ELISA, Mann-Whitney U test, n=10 **B)** T-cell cytokine production following 7 days stimulation measured by ELISA, Mann-Whitney U test, n=10

### Long-term effect of BCG on response to Influenza and SARS-CoV-2 stimulation of PBMCs from Indian and European volunteers

As a live-attenuated vaccine, BCG is known to induce long-term effects on the innate immune system and provide non-specific protection against various infections (11). This phenomenon has been termed trained immunity and is functionally characterised by augmented responsiveness of innate immune cells (12). Given that Indian and European individuals have significantly different chromatin accessibilities and differential transcriptional and functional responses to viral stimulation, we hypothesised that they would also have distinct responses to BCG vaccination. Therefore, we analysed viral response of PBMCs of Indian and European individuals before and 10-12 weeks after BCG vaccination.

Transcriptional analysis of unstimulated PBMCs before and after BCG showed minimal differences in the Indian group, whereas no differences were observed in the European group (**Figure 4A**). On the other hand, viral stimulation induced significant changes in the transcriptome before and after BCG vaccination. The magnitude of response to either Influenza or SARS-CoV-2 was overall greater in the European group than in the Indian group, both before and after BCG vaccination (**Figure 4B**). The number of DEGs after influenza virus stimulation significantly changed (from 1307 to 1143) in Indian individuals, compared to Europeans (from 2550 to 2843, Fisher Exact test, p<0.00001).

**Figure 4.**
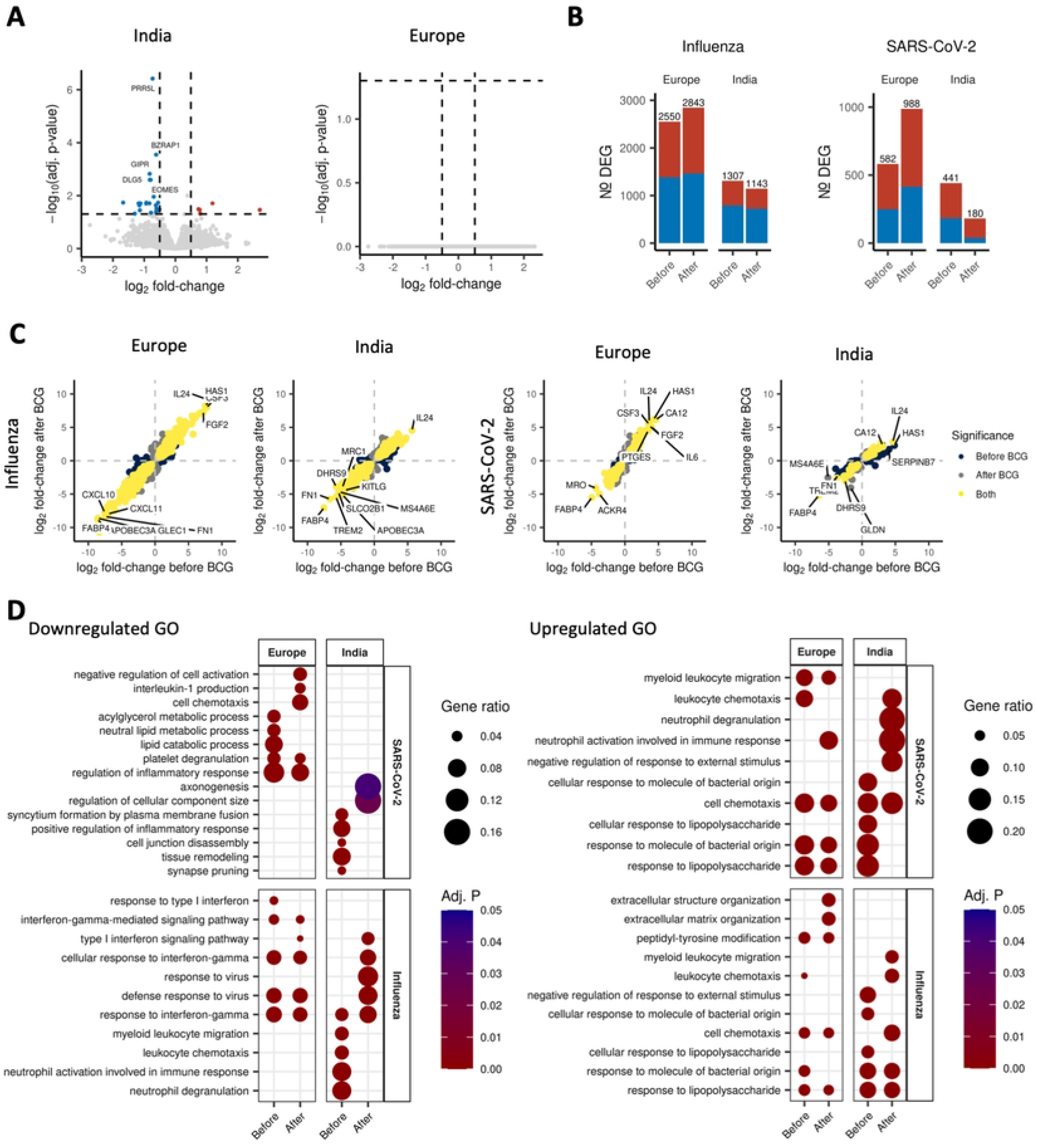
The transcriptional response to viral stimulation before and after BCG vaccination. **A)** Volcano plot showing the effect of BCG vaccination on the transcriptome (Absolute log2-transformed fold change > 0.5). **B)** Barplot indicating the number of differentially expressed genes for each viral stimulation compared to RPMI-treated PBMCs before and after BCG vaccination. Blue indicates downregulation, red indicates upregulation. **C)** Consistency of the response to viral stimulation before and after BCG. Each point represents a gene, plotted by the logFC per stimulation before and after BCG vaccination. **D)** Dotplot of GO term enrichment results showing the transcriptional modules that are regulated following stimulation, before and after BCG.

The response to SARS-CoV-2 showed an even clearer difference. In the European group, the number of DEGs increased substantially after BCG vaccination (from 582 to 988). Surprisingly, the number of DEGs strongly decreased in Indian individuals after BCG vaccination (from 441 to 180) (**Figure 4B**). Systemic comparison of the direction of regulation per gene, and the analysis revealed that most genes are regulated in a consistent fashion between the groups, although population-specific genes were also observed. This pattern was consistent for both stimulations (**Figure 4C**).

GO enrichment analysis of the differentially expressed genes upon stimulation underlined the differences in response to viral stimuli between European and Indian individuals (**Figure 4D**). These results align with the previous data showing that European and Indian individuals have different chromatin accessibility profiles related to innate and adaptive immunity, respectively. Stimulation with SARS-CoV-2 in the European group down-regulated metabolic processes such as lipid metabolism and regulation of inflammatory response before BCG. However, this shifted towards inhibition of the IL-1 production pathway and negative regulation of cell activation with the persistent presence of regulation of inflammatory response after BCG. In contrast, Indian individuals had no overlap with the active pathways in European individuals. Positive regulation of inflammatory response was downregulated by SARS-CoV-2 stimulation in cells collected before BCG; however, this disappeared after BCG vaccination. After the BCG vaccination, the two most significantly down-regulated processes in response to SARS-CoV-2 were regulation of cellular component size and axonogenesis (**Figure 4D**). There was an emphasis on the induction of myeloid leukocyte migration and leukocyte chemotaxis in European individuals before BCG, which was absent in Indian individuals. Interestingly, they had a very distinct response to SARS-CoV-2 after BCG. While the vaccination boosted the expression of genes important for neutrophil function in Indian individuals, it did not change the balanced distribution of different immunological response pathways in European individuals.

Furthermore, down-regulated pathways related to IFNγ pathways in response to influenza did not diverge in European individuals before and after BCG. On the other hand, in the case of Indian individuals, neutrophil degranulation and response to IFNγ pathways were down-regulated before BCG, while defense response to virus-associated pathways was down-regulated after the vaccination. When we checked upregulated pathways, there was no specific significant pattern for European individuals before and after BCG.

After analysing the effect of BCG on transcriptional profiles of both groups of individuals, we sought to understand the functional outcome of the process. We measured pro-inflammatory cytokines (IL-6, TNFα, IL-1β) and IFNγ from stimulated PBMCs of both populations before and after BCG (**Figure 5**). As we presented before in the transcriptional data with higher innate immune activation in the Indian group after BCG, pro-inflammatory cytokine secretion was elevated in these individuals after vaccination in response to Influenza and SARS-CoV-2. On the other hand, in European individuals we were not able to identify significant changes in innate immune responses after vaccination, most likely due to the large variation. This pattern was similar for IFNγ production, but with a significant increase in the Indian group following vaccination in response to SARS-CoV-2.

**Figure 5.**
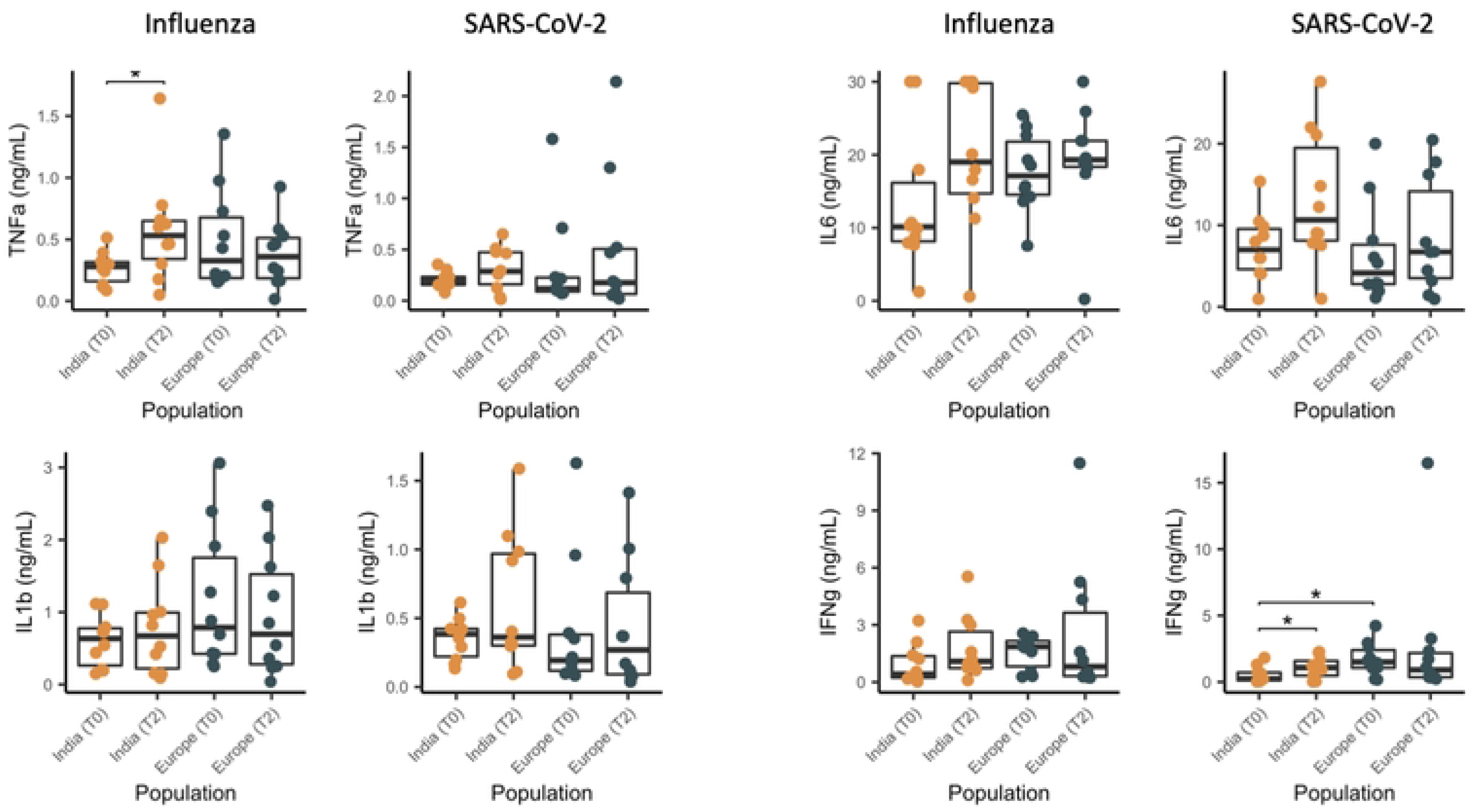
Indian and European individuals display distinctive differences in cytokine production before and after BCG vaccination. Cytokine production increases more prominently in Indian individuals compared to European individuals after BCG vaccination. T0: Baseline, T2: 10-12 weeks post-vaccination, n=10. Wilcoxon paired test used to compare within groups, Mann-Whitney U test used to compare between groups.

## Discussion

In the present study, we explored the differences in antiviral immune responses between individuals of European and Indian origin before and after BCG vaccination. Our analyses show divergent viral responses between populations. Identical transcriptional modules are activated upon stimulation, yet the extent of activation is greater in Europeans, while a more regulated tolerant response was observed in Indians. Vaccination with BCG, that has been earlier proposed to induce heterologous protective anti-viral responses (11), also induced different effects in the two populations: interestingly, the overall transcriptional responses were larger in cells from European donors after ex-vivo stimulation with SARS-CoV-2 or influenza viruses, while becoming more regulated and limited in Indians. This resulted in a more focused and likely more effective functional immune response against viruses in individuals of Indian origin.

There are several hypotheses for the different impact of COVID-19 observed in Indian and European populations. First, Indians likely experience more intense infectious pressure compared to Europeans. This may lead to a more activated state of the innate immune response during homeostasis, which potentially ‘primes’ the innate immune cells to respond more adequately to stimulation, a process termed trained immunity (12). Furthermore, constant infectious pressure may lead to a more developed adaptive immune system, and consequently a more controlled immune response to viral challenges. Together, the cumulative effect of all past vaccinations, infections and other viral challenges substantially influence the present human immune response (13). Secondly, substantial differences exist in the genetic background between the two populations. While we did not consider the genetic aspect in the current study, previous work set out to map the genetic influence of viral infection (14) and cytokine production (15) between genetically divergent populations. In a recent study comparing anti-viral immune responses between individuals of European and African ancestry, the individuals with a high European ancestry had an immune response associated with an increased type I interferon response upon influenza stimulation, compared with the African population (14). This could confirm our results, although the comparison was between different populations. Third, differences in cross-reactive T-cell populations may provide a degree of adaptive immunity to Influenza or SARS-CoV-2. Indeed, studies have shown that sub-Saharan African populations had higher serological cross-reactivity to SARS-CoV-2, even before the start of the pandemic (5). Altogether, both host and environmental factors may determine the differences between the basal chromatin accessibility and anti-viral responses in Europeans and Indians, and future studies are warranted to investigate this.

BCG vaccination has been shown to induce beneficial long-term effects on the innate arm of the immune system, and provide non-specific protection against various infections (11). A recent large phase 3 BCG trial in elderly Europeans failed to show a significant difference in the total number of infections between vaccinated and non-vaccinated individuals (16), although a positive effect on mortality has been suggested (17). In contrast, initial information suggests positive effects of BCG vaccination in a phase 3 clinical trial of elderly individuals in India (18,19). Our data suggests that such positive effects may be related to a more tolerant but effective anti-SARS-CoV-2 immune response in Indians after vaccination with BCG, which would avoid hyperinflammation and subsequent severe forms of the disease.

There are several limitations to be considered. Firstly, all Indian individuals receive the BCG vaccination at birth, which is not administered to the Dutch participants. Therefore, the current study shows the effect of BCG re-vaccination in the Indians, rather than their first exposure to the vaccine. Secondly, despite the clear and statistically significant differences we observe here, the current study is limited by the number of participants. We did not have access to more biological samples from volunteers in India due to logistical aspects related to the pandemic itself. At later times during the pandemic, when the accessibility improved, the vaccination programs with specific anti-COVID19 vaccines needed to be taken into account. Further studies, especially those investigating the genetic aspect of differential anti-viral responses between populations, should focus on larger population-based cohorts from different genetic ancestries to properly power their studies.

In summary, this study shows a significantly different anti-viral response between Indian and European individuals. Individuals of Indian ancestry display a less exuberant transcriptional response upon exposure of their immune cells to viral stimulation compared to Europeans. Furthermore, BCG re-vaccination further promoted a more tolerant but effective immune response to SARS-CoV-2 in Indian, but not European, individuals. These differences could be driven by infectious burden, previous vaccinations, lifestyle, heterologous leukocyte immune cell populations, trained immunity etc, and future studies are warranted to explore these possibilities in greater detail.

## Acknowledgements

The authors extend their gratitude to all study participants.

## Author contributions

Conceptualization and design: MGN, AV, YL

Data analysis: MZ & BG

Patient recruitment, collection of biological material and experimental work: BG, GK, PAD, MPB, JHAM, JDA

Writing – initial draft: BG, MZ, MGN, YL

Writing – review & editing: All co-authors.

## Methods

### Study design and sample collection

Indian participants are healthy healthcare workers of St. John’s Medical College Hospital, Bangalore, India, invited to participate between October 2019 and June 2021 (20). All included Indian participants received the BCG vaccination at birth. European participants were healthy, non-smoking young volunteers, undergoing BCG vaccination in the BCG-Booster study. Participants did not have a history of past BCG vaccination. Exclusion criteria were acute illness within two weeks before receiving the BCG vaccine and using medication, including non-steroidal anti-inflammatory drugs (NSAIDs), within 4 weeks (except for oral contraceptives) before vaccination. Female participants were screened for pregnancy.

### Ethics approval

This study was performed in accordance with the relevant guidelines and regulations stated in the Declaration of Helsinki. Ethical approval for the Indian participants was given by the Institutional Ethics Ethical Review Committee of St. John Medical College Hospital, Bangalore, IEC Ref no: (IEC/1/896/2018). Ethical approval for the European individuals was given by the Arnhem-Nijmegen Ethical Committee, number NL58219.091.16.

### PBMC isolation

Venous blood of the European study participants was collected into 10 mL EDTA tubes (BD Vacutainer, USA). PBMC isolation was performed using density centrifugation of blood diluted 1:1 in PBS and layered over Ficoll-Paque Plus (Cytiva, USA). Then, the PBMC layer was collected and washed twice in cold PBS. After cell counting, the PBMCs were preserved in bovine calf serum (BCS) containing 10% DMSO, incubated overnight at −80°C in Mr. FrostyTM freezing container; then stored in liquid nitrogen to be used in sequencing and stimulation assays. Blood from the Indian study participants was collected in Na-Heparin tubes (BD, Franklin Lakes NJ, USA) and diluted 1:1 with PBS (Gibco by Life Technologies, Washington, DC, USA) + 2% FBS (Gibco). PBMC isolation was performed using 15ml ACCUSPIN tubes (Sigma-Aldrich) by density centriguation following the manufacturer’s infstructions. PBMCs from the buffy coat were washed twice with PBS + 2% FBS, then re-suspended at 10 × 10^6^ cells/mL in cryopreservation medium (90% FBS and 10% DMSO), incubated overnight at −80°C (in Mr. FrostyTM freezing container; Nalgene, Rochester, New York, U.S.) and were stored in liquid nitrogen until further analyses.

### Bulk ATAC sequencing

Untreated PBMCs were subjected to tagmentation to prepare for ATACseq. The cells were lysed with TDE1 (Tagment DNA enzyme, Illumina, cat# 20034197). After lysis, DNA fragments were eluted following Qiagen Min Elute kit protocol. DNA was subsequently PCR amplified by using KAPA HiFi Hotstart Ready Mix (KAPA Biosystems) and Nextera Index Kit (Illumina) primers followed by reverse phase 0.65 x SPRI beads purification and a QIAquick Spin Column (QIAGEN) purification. Amplified DNA libraries were sequenced with an Illumina NextSeq 500 at a read length of 38 bp.

### PBMC stimulation experiments

The PBMCs were thawed and washed with 10mL Dutch modified RPMI 1640 medium (Roswell Park Memorial Institute; Invitrogen, USA, cat # 22409031) containing 50 µg/mL Gentamicine (Centrafarm, The Netherlands), 1 mM Sodium-Pyruvate (Thermo Fisher Scientific, USA, cat #11360088), 2 mM Glutamax (Thermo Fisher Scientific, USA, cat #35050087) supplemented with 10% Bovine Calf Serum (Fisher Scientific, USA, cat #11551831) twice. Afterwards, cells were counted via Sysmex XN-450. PBMCs (4×10^5^ cells/well) were laid in sterile round bottom 96-well tissue culture treated plates (VWR, The Netherlands, cat #734-2184) in Dutch modified RPMI 1640 medium containing 50 µg/mL Gentamicine, 1 mM Sodium-Pyruvate, 2 mM Glutamax supplemented with 10% human pooled serum. The cells were rested for an hour in a 37°C incubator supplied with 5% CO_2_ before stimulation. Following, stimulations were done with two heat-inactivated viral stimuli; SARS-CoV-2 Wuhan Hu-1 strain (2.8×10^3^ TCID50/mL), Influenza California A (3.6×10^3^ TCID50/mL). The PBMCs were incubated with the stimulants for 24 hours to detect IL-1β, TNF-α, IL-6, IL-1Ra, and 7 days to detect IFN-γ, IL-17 and IL-10. Supernatants were collected and stored in −20°C.

### Bulk RNA sequencing

PBMCs were lysed with Qiagen RBP buffer following 24 hours simulation and stored at −80 until isolation. RNA was isolated using Qiagen RNeasy kit following manufacturer’s instructions. A total amount of 200ng RNA per sample was used for the preparation of RNA sequencing libraries using the KAPA RNA HyperPrep Kit with RiboErase (HMR) (KAPA Biosystems). In short, oligo hybridization and rRNA depletion, rRNA depletion cleanup, Dnase digestion, Dnase digestion cleanup, and RNA elution were performed according to protocol. Fragmentation and priming were performed at 94 C for 6:30 min. First strand synthesis, second strand synthesis, and A- tailing was performed according to protocol. For the adapter ligation, a 7 μM stock was used (NextFlex DNA barcodes, Bioo Scientific). First, and second post-ligation cleanup was performed according protocol. For the library amplification, 6 cycli were used. The library amplification cleanup was performed using a 0.8x bead-based cleanup. Library size was determined using the High Sensitivity DNA bioanalyzer (Agilent Technologies), library concentration was measured using the DeNovix dsDNA High Sensitivity Assay (DeNovix). Sequencing was performed using an Illumina NextSeq 500, 38-bp paired-end reads were generated.

### Quantification and statistical analysis

#### Cytokine production upon stimulation

Secreted cytokine levels from supernatants following stimulation were quantified by ELISA (IL-1β cat # DLB50, TNF-α cat # STA00D, IL-6 cat # D6050, IL-1Ra cat # DRA00B, IFN-γ cat #DY285B, IL-10 cat #DY21YB, IL-17 cat# DY317, R&D Systems, USA), following the instructions of the manufacturer. Mann-Whitney U test was used to compare cytokine levels between populations, and the Wilcoxon paired test to compare within populations.

#### Bulk ATAC sequencing

ATAC sequencing reads were pre-processed using the nfcore/atacseq pipeline ((21), v1.2.1) implemented in Nextflow ((22), v21.04.3) using default setting and the human GRCh39 genome. We considered the consensus peak set called by MACS2 in broad mode. Peaks located on sex chromosomes were removed prior to further analysis, and remaining peaks were annotated using HOMER (23). Differential peak abundance to compare the populations was performed using linear models in DESeq2 ((24), v1.30.1), incorporating individual’s age and sex in the model. Resulting p-values were corrected across all peaks using Benjamini-Hochberg, and adjusted P-values < 0.05 were considered significant.

#### Bulk RNA sequencing

Bulk RNA sequencing reads were pre-processed using the publicly available nfcore/rnaseq pipeline ((21), v2.0) using default settings and the human GRCH38 genome. Differential gene expression was implemented using linear models in DESeq2 ((24), v1.30.1). Models included individual’s sex to estimate the BCG vaccination effect and baseline differences between the populations. To compare each viral stimulation against RPMI-treated control samples, a paired design was used that included each individual’s study ID. Genes with total read counts < 20 were removed prior to further analysis. Resulting p-values were corrected across all genes using Benjamini-Hochberg, and adjusted P-values < 0.05 were considered significant.

## Resource availability

Further information and requests for resources and reagents should be directed and will be fulfilled by the lead contact, Mihai Netea

## Data availability

Bulk transcriptomics and bulk ATAC are freely available and have been deposited at the European Genome-phenome Archive (EGA), which is hosted by the EBI and CRG, under accession number EGAS00001006417.

## Code availability

Codes used in this study are freely available on Github (https://github.com/CiiM-Bioinformatics-group/INDIA.git).

